# Critical assessment of missense variant effect predictors on disease-relevant variant data

**DOI:** 10.1101/2024.06.06.597828

**Authors:** Ruchir Rastogi, Ryan Chung, Sindy Li, Chang Li, Kyoungyeul Lee, Junwoo Woo, Dong-Wook Kim, Changwon Keum, Giulia Babbi, Pier Luigi Martelli, Castrense Savojardo, Rita Casadio, Kirsley Chennen, Thomas Weber, Olivier Poch, François Ancien, Gabriel Cia, Fabrizio Pucci, Daniele Raimondi, Wim Vranken, Marianne Rooman, Céline Marquet, Tobias Olenyi, Burkhard Rost, Gaia Andreoletti, Akash Kamandula, Yisu Peng, Constantina Bakolitsa, Matthew Mort, David N. Cooper, Timothy Bergquist, Vikas Pejaver, Xiaoming Liu, Predrag Radivojac, Steven E. Brenner, Nilah M. Ioannidis

## Abstract

Regular, systematic, and independent assessment of computational tools used to predict the pathogenicity of missense variants is necessary to evaluate their clinical and research utility and suggest directions for future improvement. Here, as part of the sixth edition of the Critical Assessment of Genome Interpretation (CAGI) challenge, we assess missense variant effect predictors (or variant impact predictors) on an evaluation dataset of rare missense variants from disease-relevant databases. Our assessment evaluates predictors submitted to the CAGI6 Annotate-All-Missense challenge, predictors commonly used by the clinical genetics community, and recently developed deep learning methods for variant effect prediction. To explore a variety of settings that are relevant for different clinical and research applications, we assess performance within different subsets of the evaluation data and within high-specificity and high-sensitivity regimes. We find strong performance of many predictors across multiple settings. Meta-predictors tend to outperform their constituent individual predictors; however, several individual predictors have performance similar to that of commonly used meta-predictors. The relative performance of predictors differs in high-specificity and high-sensitivity regimes, suggesting that different methods may be best suited to different use cases. We also characterize two potential sources of bias. Predictors that incorporate allele frequency as a predictive feature tend to have reduced performance when distinguishing pathogenic variants from very rare benign variants, and predictors supervised on pathogenicity labels from curated variant databases often learn label imbalances within genes. Overall, we find notable advances over the oldest and most cited missense variant effect predictors and continued improvements among the most recently developed tools, and the CAGI Annotate-All-Missense challenge (also termed the Missense Marathon) will continue to assess state-of-the-art methods as the field progresses. Together, our results help illuminate the current clinical and research utility of missense variant effect predictors and identify potential areas for future development.

## Introduction

Predicting the significance of genetic variation is an ongoing challenge that is essential for determining genetic susceptibility to disease and identifying causal variants in rare disease diagnosis [1]. Clinical sequencing laboratories often struggle with the interpretation of rare and *de novo* variants seen in patients, classifying them as variants of uncertain significance (VUS) due to a lack of available evidence about their pathogenicity. Interpretation of missense variants is of particular interest due to their frequent occurrence and wide range of potential effects on protein function and clinical phenotypes, ranging from no effect to either an adaptive effect or a highly penetrant pathogenic loss or gain of function [2]. The American College of Medical Genetics and Genomics (ACMG) and the Association for Molecular Pathology (AMP) have developed guidelines for clinical variant interpretation that describe how to integrate numerous lines of evidence, including predictions from computational tools, when classifying a variant as pathogenic or benign [3], and ClinGen has issued an updated clinical recommendation for the use of computational tools [4].

In the last two decades, many computational tools have emerged to predict the pathogenicity or clinical significance of missense variants, leveraging variant and gene-level annotations such as evolutionary conservation and protein structural properties as predictive features [5, 6]. These methods are called variant effect predictors or variant impact predictors. Predictions from individual tools often disagree, which motivated the development of ensemble methods, or meta-predictors, trained to aggregate predictions from multiple tools. Meta-predictors tend to have improved performance over their component predictors, but rely on the continued development of individual predictors that incorporate information from new or complementary predictive features. Therefore, it is important to assess the performance of both meta-predictors and individual predictors on the task of missense variant pathogenicity prediction.

The goal of the CAGI Annotate-All-Missense challenge, also termed the Missense Marathon, is to conduct an ongoing assessment of missense pathogenicity predictors (both existing tools and those newly submitted to the CAGI challenge) using variants that have been classified as pathogenic or benign in clinical variant databases or identified as disease-causing variants since the close of the most recent challenge. Teams submitting to the challenge were asked to provide prediction scores for all possible missense single nucleotide variants (SNVs) in the human reference genome, based on dbNSFP v4 [7], similar to existing missense variant effect predictors with precomputed scores. A preliminary, limited assessment of missense predictors was previously performed as part of CAGI5 [1]. Here we perform a more extensive analysis of missense variant effect predictors for the CAGI6 challenge, using variants with pathogenicity information made available between November 2021 and April 2023 in the evaluation set.

## Results

### Evaluation dataset

We curated a dataset of rare (allele frequency *<* 0.05) missense variants classified in ClinVar [8] as either Pathogenic or Benign with at least one star, or listed as disease-causing (DM) entries in the Human Gene Mutation Database (HGMD) [9]. Our dataset was restricted to variants that were newly added to these databases after the close of the CAGI6 Annotate-All-Missense challenge in October 2021. We consider the ClinVar and HGMD data both together and separately in the analyses below, to explore differences between the databases. Variants with pathogenicity information available in ClinVar, HGMD, or UniProt [10] prior to the close of the challenge were explicitly excluded. Common variants were removed, as they should be considered benign per the updated standalone BA1 rule in the ACMG/AMP guidelines [11]. Additional details of dataset construction are provided in Methods. The resulting dataset contained 6,103 pathogenic and 4,353 benign variants from 2,115 genes, with an allele frequency distribution shown in Fig. S1.

### Missense variant effect predictors

We evaluated the performance of 60 missense variant effect predictors, of which 12 were submitted by 6 teams to the CAGI6 Annotate-All-Missense challenge. The additional tested methods include predictors commonly used by the clinical genetics community and recently developed deep learning methods for missense variant interpretation (listed in Table S1 and Methods). All assessed predictors were either released before the close of the challenge or did not train on variant pathogenicity data released after the challenge ended, to ensure no overlap with the evaluation set. Nonetheless, other more subtle forms of circularity may exist (e.g. [12]) and are discussed in more detail below. We also note that some predictors were trained or fine-tuned on variants from population databases such as gnomAD [13], which contain allele frequency information but not clinical classifications; therefore, we did not specifically exclude such variants from the evaluation set.

For presentation clarity, in most analyses, we show results for a select subset of 26 predictors: the top-performing model from each team from the CAGI6 challenge, predictors widely used by the clinical genetics community, and recently developed methods that have garnered interest. We note that 5 of the 6 top-performing CAGI6 team submissions are (nearly) identical to previously published methods—3Cnet [14], MetaRNN [15], MISTIC [16], SNPs&GO [17], and VESPAl [18]. In figures, we label them with both their familiar method name and the submitting team identifier (e.g. 3Cnet/3billion). The sixth team submission, labeled as DEOGEN2/(IB)2, uses DEOGEN2 [19] scores when available and rescaled PROVEAN [20] scores when not. Of the remaining 20 highlighted predictors, 15 are commonly used in clinical genomics and research applications: BayesDel (with and without allele frequency) [21], CADD [22], ClinPred [23], Eigen [24], FATHMM-XF [25], MutationAssessor [26], MutPred2 [27], M-CAP [28], PolyPhen2 [29], phyloP [30], PROVEAN [20], REVEL [31], SIFT4G [32], and VEST4 [33]. Among the 5 recently developed methods (AlphaMissense [34], ESM-1b [35], EVE [36], PrimateAI-3D [37], and VARITY [38]), all but VARITY are deep learning methods that are not supervised on known variant pathogenicity labels. Correlations between the predictions from these tools, as measured on our evaluation dataset, are shown in Fig. S2.

In addition to these 26 predictors shown in the figures, summary metrics for the full set of 60 tested predictors are provided in Table S1.

### Full ROC curve performance

For each predictor, we first constructed its Receiver Operating Characteristic (ROC) curve based on the full evaluation dataset and computed the Area Under the ROC (AUROC) (Fig. 1 and Table S1). Note that we further break down performance within different subsets of the dataset and within different regions of the ROC curve below. We also show performance for meta-predictors and non-meta-predictors (individual predictors) separately in all figures, since we find that meta-predictors tend to achieve higher AUROCs by combining scores from individual predictors, consistent with previous observations. We also distinguish predictors that incorporate allele frequency as an explicit predictive feature, due to limitations in their use with other ACMG/AMP lines of evidence.

**Fig. 1:**
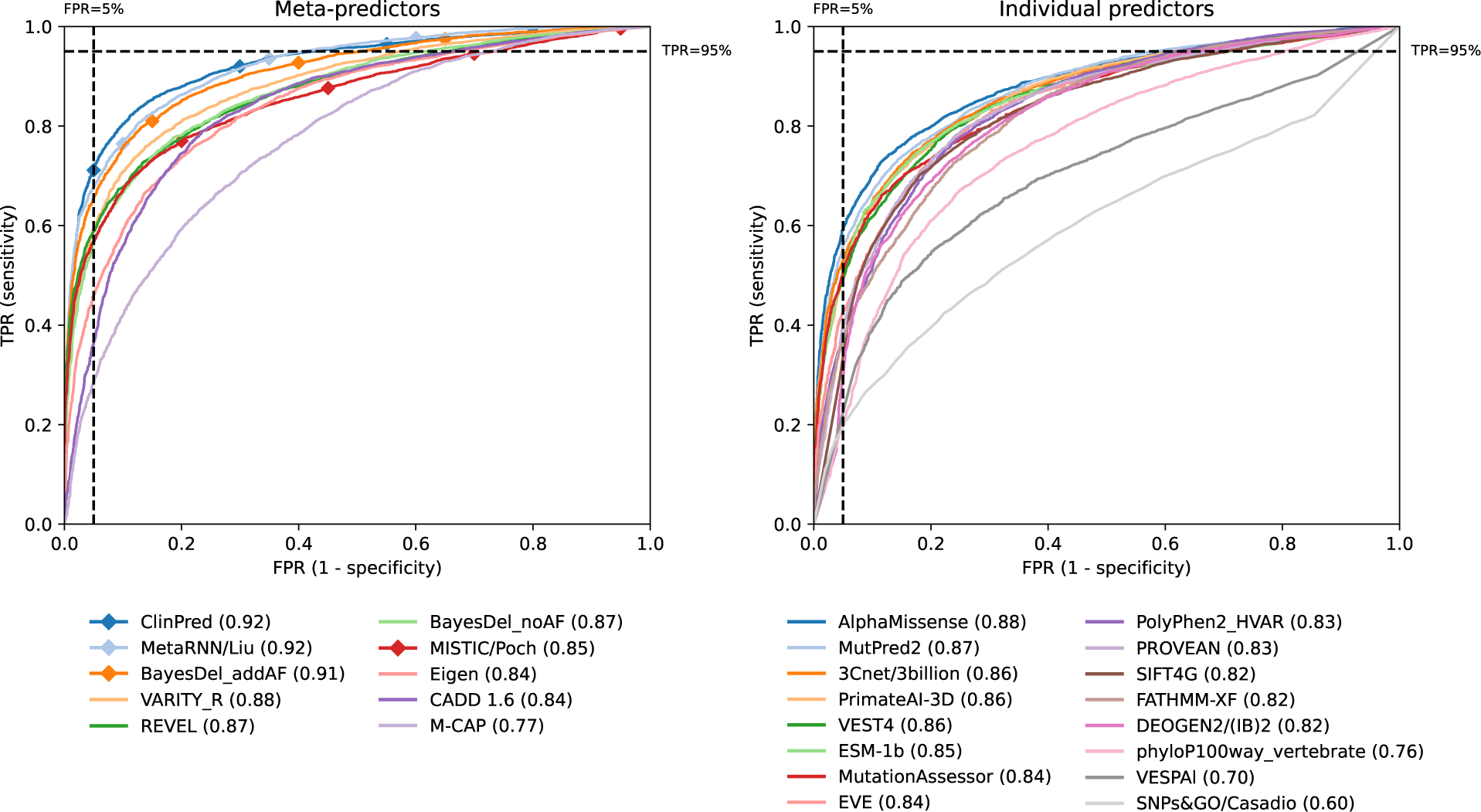
Full ROC curve performance. We show the ROC curves and AUROCs for meta-predictors (left) and individual predictors (right) on the full evaluation dataset. Predictors marked by diamonds use allele frequency as a feature. The black dashed lines at 5% FPR and 95% TPR demarcate the boundaries of the high-specificity and high-sensitivity regions, respectively, which are enlarged in Fig. 2.

The meta-predictors with the highest AUROC on the full evaluation dataset reach an AUROC of 0.92 (ClinPred, MetaRNN) for methods that explicitly include allele frequency as a predictive feature (marked with diamonds in Fig. 1), and an AUROC of 0.88 (VARITY_R) for methods that do not explicitly include allele frequency. Although we limited the evaluation dataset to rare variants, the methods that explicitly use allele frequency likely benefit from the remaining allele frequency imbalance in the evaluation dataset, in which pathogenic variants have lower allele frequencies than benign variants (Fig. S1). We further explore the effect of allele frequency below. The individual predictor with the highest AUROC on the full evaluation dataset (AlphaMissense) also reaches an AUROC of 0.88. In general, there are multiple predictors with AUROCs within a few percentage points of one another—including AlphaMissense, MutPred2, 3Cnet, PrimateAI-3D, and VEST4—indicating that a number of approaches all have strong performance. Among the deep learning methods that do not supervise on labeled pathogenic or disease-causing variants, those that model both protein structure and protein language (AlphaMissense and PrimateAI-3D) slightly outperform those that model only protein language (ESM-1b and EVE).

The above results were computed for each predictor on only the subset of variants from the evaluation dataset that were scored by that predictor, ignoring missing predictions. However, 8 out of 26 predictors do not report scores for at least 5% of the evaluation dataset (Fig. S3). Most notably, EVE does not supply predictions for 37% of the dataset. Therefore, in Fig. S4, we compare performance on the full dataset (*n* = 10, 456) to performance on the smaller set of variants that are scored by all predictors (*n* = 4, 769). 23 out of 26 predictors have higher performance on the latter set, suggesting that variants that are not scored by some predictors tend to be harder to predict. We also note a depletion of benign variants relative to pathogenic variants in the smaller set of variants scored by all predictors. However, we find that the ordering of predictors by AUROC is largely similar for both sets of variants.

### ClinVar and HGMD data subsets

Pathogenic variants in our evaluation dataset were sourced from two databases with very different curation strategies. ClinVar is a database of public variant classifications primarily submitted by genetic testing laboratories and classified using ACMG/AMP criteria, whereas HGMD is a licensed database that collates disease-relevant variants from the primary literature, including basic research studies. To distinguish performance on the two databases, we constructed two subsets of our evaluation dataset: one containing pathogenic variants only from ClinVar, and the other containing pathogenic variants only from HGMD. In both cases, all benign variants were from ClinVar, since HGMD does not curate benign variants. Fig. S5 shows the difference in performance on these two data subsets. All predictors have higher performance on the evaluation subset containing pathogenic variants from ClinVar, suggesting that there is a qualitative difference between pathogenic variants obtained from the two databases. However, predictor rankings are largely similar in the two cases.

### High-specificity and high-sensitivity performance

The AUROC metric used above aggregates performance across all possible decision rules, or score thresholds, for separating predicted benign variants from predicted pathogenic variants. However, in practice, only a single decision rule is typically used for a particular application. While not all practitioners may choose the same decision rule, there are two general regimes in which computational predictors of missense variant pathogenicity are most likely to be used. First, for clinical variant interpretation, high confidence classifications of pathogenicity are required when reporting results to patients. Especially when reporting secondary genomic findings, which are putatively pathogenic variants of concern unrelated to the original reason for testing [39], false positives should be minimized to avoid overdiagnosis. Accordingly, practitioners will employ a decision rule with a low false positive rate (FPR), or equivalently, high specificity. To measure performance in this setting, we examined the high-specificity region (FPR *≤* 5%) of the ROC curves from Fig. 1 (Fig. 2a). (This particular 5% FPR threshold is arbitrary but represents a useful decision rule in this scenario.) Second, for exploratory analysis of whole-exome or whole-genome sequencing data, high sensitivity is often desired. For example, in a research environment, when analyzing data from a patient with an undiagnosed genetic disorder, computational predictors can be used to narrow down a list of VUS to those variants that should be prioritized in follow-up studies, ideally without mistaking the true pathogenic variant as benign. Accordingly, practitioners will employ a decision rule with a high true positive rate (TPR), or equivalently, high sensitivity [40]. To measure performance in this setting, we examine the high-sensitivity region (TPR *≥* 95%) of the ROC curves (Fig. 2b). For both the high-specificity (FPR *≤* 5%) and high-sensitivity (TPR *≥* 95%) regions of the ROC curves, we compute a normalized area under the curve in these regions [41]. Table S1 lists the full-curve AUROC, high-specificity AUROC, and high-sensitivity AUROC for all 60 predictors included in our evaluation.

**Fig. 2:**
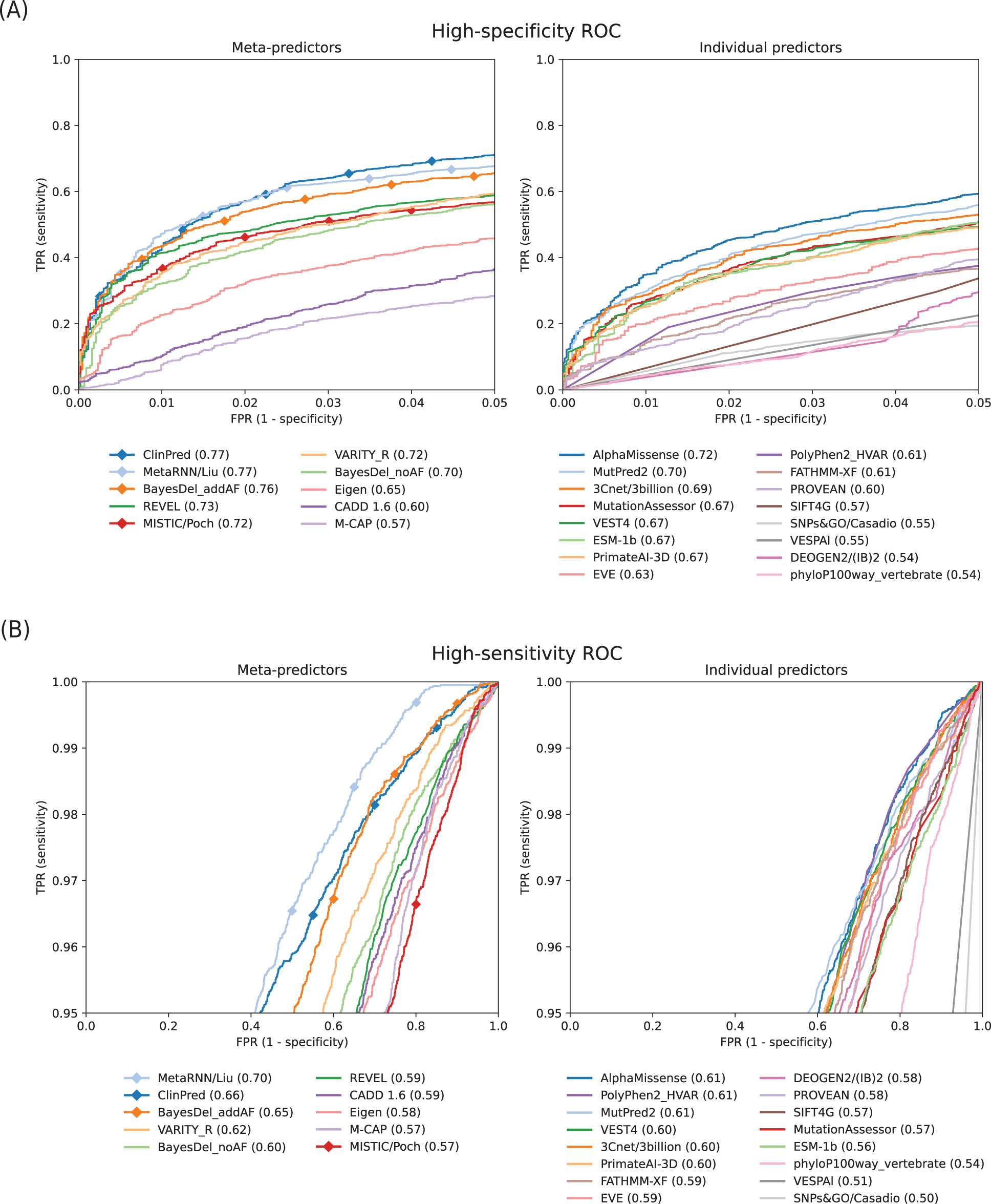
Performance in high-specificity and high-sensitivity regimes. We show enlarged portions of the ROC curves from Fig. 1 to focus on (A) the high-specificity region (FPR *≤* 5%) and (B) the high-sensitivity region (TPR *≥* 95%) for meta-predictors (left) and individual predictors (right). We also show the normalized area under the curve in these regions (normalized such that a perfect classifier gets a score of 1 and a random classifier gets a score of 0.5). Predictors marked by diamonds use allele frequency as a feature.

Notably, the performance of some predictors varies substantially between the two classification regimes. MetaRNN, which uses allele frequency as a predictive feature, excels in the high-sensitivity region, particularly for true positive rates that approach 100%. On the other hand, MISTIC performs well in high-specificity regions but struggles in high-sensitivity regions, indicating its suitability for clinical variant classification rather than exploratory research. Among the individual predictors, PolyPhen2_HVAR has strong performance in the high-sensitivity region, but lower performance relative to other predictors in the high-specificity region, whereas MutationAssessor and ESM-1b both have lower performance in the high-sensitivity region. These findings underscore the notion that different methods may be better suited for different clinical or research applications.

### Effect of allele frequency

As illustrated in Fig. S1, pathogenic variants tend to have lower allele frequencies than benign variants in our evaluation dataset. This trend is expected, as deleterious variants are more likely to be under negative selection and therefore less common in the human population. However, for many clinical use cases, it is important to be able to distinguish very rare benign variants from pathogenic variants, for example in data from rare disease patients. Methods that utilize allele frequency as a predictive feature might struggle with variant classification in this setting. To evaluate the effect of allele frequency on performance, we binned the benign variants in our evaluation dataset by their allele frequencies and compared performance when differentiating benign variants in each bin from the full set of pathogenic variants (Fig. 3). Three of the four methods that utilize allele frequency as a predictive feature—ClinPred, MetaRNN, and BayesDel_addAF, which are also top-performing predictors in the above analyses—show a marked performance decrease on very rare benign variants. Despite this decline in performance, MetaRNN and ClinPred still outperform most other predictors in distinguishing very rare benign variants from pathogenic variants, indicating that their predictions are not excessively reliant on allele frequency. VARITY_R also has notably high performance on very rare benign variants. The effect of including allele frequency as a predictive feature can also be illustrated by comparing the two versions of BayesDel with and without allele frequency. BayesDel_addAF (which performs 4% better than BayesDel_noAF on the full-dataset AUROC) has much higher performance in most of the benign allele frequency bins, but BayesDel_noAF outperforms BayesDel_addAF in the lowest allele frequency bin.

**Fig. 3:**
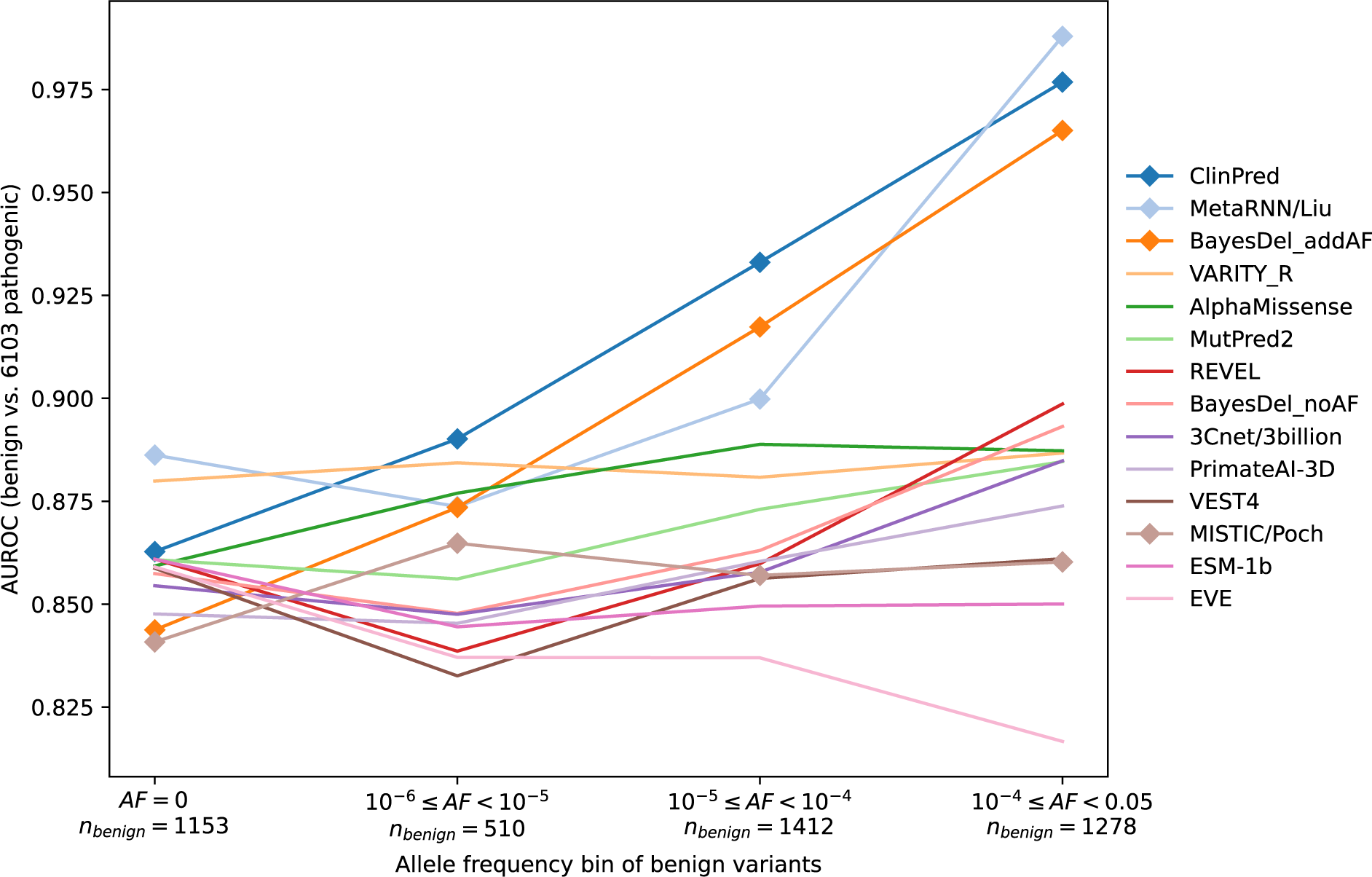
Allele frequency bias. Top-performing predictors are evaluated for distinguishing benign variants in different allele frequency bins from pathogenic variants. All 6,216 pathogenic variants were used in each evaluation, and benign variants were stratified by their allele frequencies obtained from the control cohort exomes in gnomAD v2.1.1 [13]. Predictors marked by diamonds use allele frequency as a feature.

To minimize the effect of allele frequency on our performance metrics, we created a subset of the evaluation dataset in which allele frequencies were matched between pathogenic and benign variants. We then compared performance on the full dataset to the allele frequency matched dataset (Fig. S6). Methods that use allele frequency as a predictive feature have the largest drop in performance. Some methods that do not explicitly use allele frequency also have slightly lower performance on the allele frequency matched dataset, likely because allele frequency correlates with other features used by these tools (e.g. conservation scores).

### Effect of gene label imbalance

Databases of disease-relevant variants, such as ClinVar and HGMD, have large imbalances in the ratio of pathogenic to benign variants per gene, which may reflect bias in which variants have been studied rather than the true fitness landscape for those genes [12]. To gauge the degree of label imbalance in our evaluation dataset, we tested a simple baseline model, similar to the one outlined in [34]. The baseline model assigns the same score to all variants in a gene, equal to the fraction of high-confidence missense variants from ClinVar and HGMD that were available before the cutoff date for our evaluation dataset and were labeled as pathogenic or disease-causing in those databases. On our evaluation dataset, this simple model achieves an AUROC of 0.74 (Fig. S7), rivaling the performance of the best conservation score, phyloP100way_vertebrate.

To minimize the effect of gene label imbalance on our performance metrics, we created a subset of the evaluation dataset containing an equal number of pathogenic and benign variants per gene. We then compared performance on the original dataset to the gene label-balanced dataset (Fig. 4). Many of the tested predictors, particularly many of the meta-predictors, have lower performance on the label-balanced dataset. However, predictors that do not train on labeled pathogenic or disease-causing variants (including but not limited to AlphaMissense, PrimateAI-3D, CADD, Eigen, EVE, ESM-1b, phyloP, and VESPAl) do not show a degradation in performance. The largest increase in performance on the gene label-balanced dataset is observed for SNPs&GO.

**Fig. 4:**
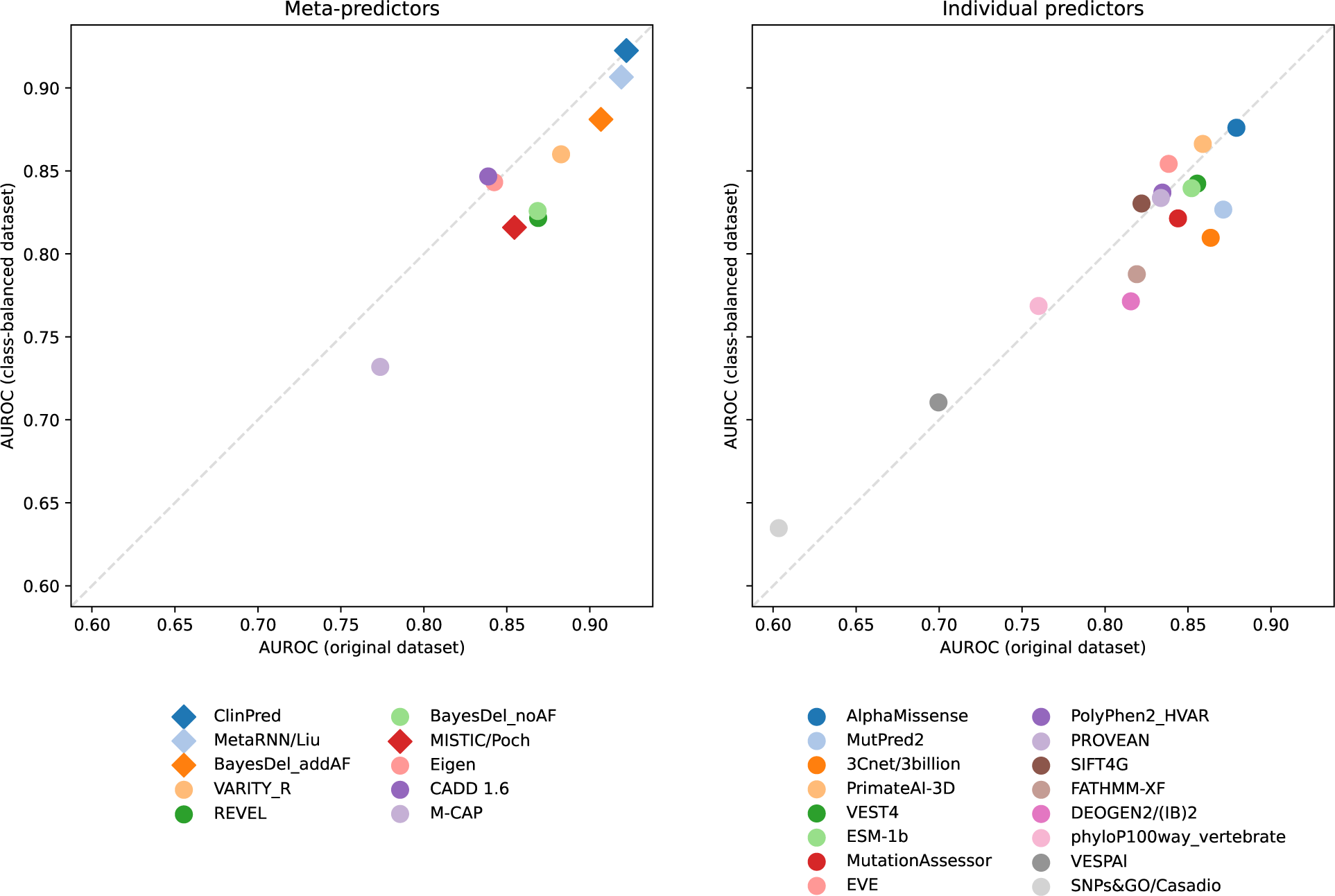
Gene label balancing. We constructed a gene label-balanced subset of our evaluation dataset containing an equal number of pathogenic and benign variants per gene. This label-balanced dataset consists of 2,140 variants from 504 genes. Performance on the label-balanced dataset (y-axis) is compared to performance on the full dataset from Fig. 1 (x-axis) for meta-predictors (left) and individual predictors (right). Predictors marked by diamonds use allele frequency as a feature.

### Effect of prior pathogenicity probability on evidence thresholds

A recently developed calibration method adopts a principled probabilistic approach to determine, for any given predictor, the thresholds at which its scores meet ACMG/AMP evidence strengths (supporting to very strong) for both pathogenicity and benignity [4, 42] using an estimated prior probability of pathogenicity [43]. For different applications, particularly in research settings, a variety of prior probabilities may be relevant. Therefore, we applied this calibration method to all tested predictors at five different prior probabilities of pathogenicity (0.02, 0.04, 0.06, 0.08, and 0.10) using our evaluation dataset. Figs. S8 and S9 display the dependence of the resulting score thresholds on the prior probability for meta-predictors and individual predictors, respectively. In all cases, lower prior pathogenicity probabilities lead to reduced evidence strengths for both benignity and pathogenicity. We note that due to the previously observed effect of allele frequency on the performance of tools that explicitly include allele frequency as a predictive feature, in future studies such tools should be calibrated separately on variants within each allele frequency bin. We also note that these tools are limited in their ability to be combined with other allele frequency-based lines of evidence, such as BA1, in the ACMG/AMP guidelines.

## Discussion

We present the first full assessment of the ongoing CAGI Annotate-All-Missense challenge, evaluating the ability of computational variant effect prediction tools to classify missense variants as pathogenic or benign under a variety of evaluation conditions. In general, we find strong performance of many predictors on an evaluation dataset of missense variants that were classified in ClinVar or added as disease-causing in HGMD after the close of the CAGI6 challenge in October 2021. Predictor rankings are largely similar when evaluated on pathogenic variants from the two databases separately.

Rather than using a single overall performance metric, it is important to evaluate missense variant effect predictors in a variety of settings that are relevant to different clinical or research applications. We examined performance in high-specificity and high-sensitivity settings separately, since high specificity is most relevant to clinical variant classification and high sensitivity is most relevant to exploratory analysis of whole-genome or whole-exome sequencing data in a research setting. We also examined performance on subsets of the evaluation dataset that were either matched for pathogenic and benign allele-frequency distributions or that included only very rare benign variants. These evaluation settings are important for applications that already use allele frequency as a separate criterion for establishing benignity (e.g. BA1 [11]) or that aim to classify missense variants within a pool of very rare variants from rare disease patients. In general, we find that predictors with strong performance tend to perform well across multiple settings, but that the specific predictor rankings differ between settings, suggesting that different predictors may be best suited to different clinical or research applications. We recommend that practitioners consider the most relevant evaluation data subsets or the most relevant portions of ROC curves for their application and choose methods based on performance in those settings, rather than examining only the full AUROC on the full evaluation dataset.

We also evaluated performance on a subset of the evaluation dataset with equal numbers of pathogenic and benign variants per gene, due to substantial imbalance in the class labels of the available variants for many genes in the full dataset. This type of imbalance is present within clinical variant databases and leads predictors trained on variants from such databases to learn gene-level properties in addition to variant-level properties for variant classification, which tends to result in reduced performance on the gene label-balanced subset. However, interpreting performance on this subset is complicated by the fact that some gene label imbalance is reflective of true biology, such as a different tolerance to mutation for different genes, while some is due to current practices in clinical testing or bias in the amount of attention and research effort devoted to particular genes and diseases. Separating the different factors contributing to gene label imbalance is an ongoing challenge in the evaluation of missense variant effect predictors. As above, we note that while the specific predictor rankings differ on this evaluation subset, predictors with strong performance in other settings also tend to perform well in the gene label-balanced setting.

For this assessment, we used recently classified or disease-relevant variants from clinical variant databases due to their relevance for evaluating clinical utility; however, we note that this source of evaluation data has several limitations in addition to the gene label imbalance discussed above. Importantly, there are likely to be some errors in the labels provided by these databases, which limit the maximum achievable performance in our evaluation. Although we attempted to reduce such errors by using high confidence labels from each database—variants with Benign or Pathogenic labels (excluding Likely Benign and Likely Pathogenic), without conflicting interpretations, and with at least 1-star from ClinVar, and DM variants from HGMD— some incorrectly labeled variants presumably remain. To account for possible systematic differences between the two databases, we report performance on the ClinVar and HGMD subsets of pathogenic variants separately, in addition to performance on the full evaluation dataset. The substantially lower performance that we observe for all predictors on the HGMD disease-causing variants likely reflects differences in annotation practices between the two databases.

In addition, many missense variant effect predictors were trained using data from clinical variant databases, and any overlap between these training variants and the variants in the evaluation dataset would inflate the performance estimates for such predictors. To avoid overlap, we specifically excluded variants from our evaluation dataset that had pathogenicity information available in ClinVar, HGMD, or UniProt, which are the most commonly used databases for training predictors, prior to the close of the CAGI6 challenge. However, it is possible that some evaluation set variants had been studied in the literature or included in other specialized databases prior to this date, where pathogenicity labels could have been available for training. Another limitation of using clinical variant databases for evaluation is the potential for circularity if predictions from any of the tested tools were considered when making pathogenicity classifications for the variants in the evaluation dataset. Based on the Richards *et al.* 2015 [3] ACMG/AMP guidelines, computational predictions had primarily been used only as supporting evidence for classification, but recent clinical recommendations [4] identified thresholds at which predictions from certain tools provide stronger levels of evidence. Therefore, it is possible that computational predictions had some influence on the most recently classified variants, which would result in inflated performance estimates for those and related tools. To enable continued unbiased assessments of missense variant effect predictors in the future, it will be essential for clinical variant databases to document the lines of evidence used for each variant classification.

Overall, our results indicate that currently available tools for missense variant effect prediction provide a powerful line of evidence for classifying missense variants of uncertain significance. While we find that meta-predictors tend to outperform their constituent individual predictors, a number of individual predictors have performance close to that of commonly used meta-predictors, particularly meta-predictors that do not explicitly include allele frequency as a predictive feature. We note continued progress in the field relative to the oldest and most cited tools, as well as recent advancement in developing individual predictors that are not trained on variants from clinical variant databases, making them less susceptible to biases in the collection and interpretation of variant data. Several such predictors achieve strong performance in our assessment, including predictors that use only unsupervised or self-supervised training schemes. These types of predictors are promising candidates to be incorporated into future meta-predictors and combined with other complementary information related to variant pathogenicity. This ongoing CAGI challenge will continue to evaluate such developments and to assess state-of-the-art methods as the field progresses.

## Methods

### Evaluation dataset construction

We created an evaluation dataset by incorporating variants from both the April 4, 2023 version of ClinVar [8], which contains both pathogenic and benign variants, and the 2023.1 Professional version of the Human Gene Mutation Database (HGMD) [9], which contains only pathogenic variants.

We assigned molecular effects to all variants using SnpEff [44], which was configured with the Ensembl 105 gene set, and retained all single-nucleotide variants that were annotated as missense in at least one affected transcript, excluding variants that were assigned a higher impact annotation (`HIGH` impact or `splice_region_variant`) in another transcript.

To ensure that all predictors were tested on variants that they had not previously seen during training, we removed (1) all variants, except those of uncertain significance, present in the November 7, 2021 version of ClinVar, (2) all `DM` variants present in the 2021.4 version of HGMD, (3) all variants, expect those of uncertain significance, present in the 2021.4 version of the UniProt Humsavar database [10], and (4) variants in the AlphaMissense validation set (used for early stopping). These cutoff dates were chosen based on the CAGI6 challenge, which closed on October 11, 2021. Motivated by PM5 [3], we also excluded variants affecting the same codon as any of the aforementioned removed variants to minimize data leakage (Fig. S10).

Among the remaining variants, we only retained those with high-confidence pathogenicity classifications in ClinVar (either `Benign` or `Pathogenic` with 1 star or above, except those with conflicting interpretations) and high-confidence disease-causing (DM) HGMD variants. We further removed variants that can be inferred to be benign by their allele frequency in gnomAD exomes v2.1.1 [13], as per the revised BA1 criterion [11]. Specifically, we removed variants with a control global allele frequency *>* 0.05 or control continental allele frequency *>* 0.05 with at least 2000 observed alleles in any of the five major continental populations: African/African American (AFR), Latino/Admixed American (AMR), East Asian (EAS), South Asian (SAS), and non-Finnish European (NFE). Furthermore, we discarded all variants that were not present in a Mendelian disease gene. For the purposes of this study, we consider Mendelian disease genes (*n* = 3465) to be those with at least one high-confidence (as described above) pathogenic variant of any mutation class in the April 4, 2023 version of ClinVar. Lastly, we excluded variants that predictors submitting to the CAGI6 Annotate-All-Missense challenge were not asked to score. Our final dataset contains 6,103 pathogenic and 4,353 benign variants.

### Procuring predictions

We evaluated 60 missense pathogenicity predictors, none of which were trained on clinical pathogenicity data released after the CAGI6 challenge deadline.

#### CAGI6 Annotate-All-Missense submissions

Six teams submitted a total of twelve models to the challenge. Submitters were asked to provide a prediction score for a pre-specified list of missense variants throughout the genome (based on dbNSFP v4 [7]), of which our evaluation dataset is a subset. All team identities and model details were hidden until the conclusion of the analysis.

#### Predictors available in dbNSFP

We obtained predictions for 40 tools from the dbNSFP v4.2a database (released on April 6, 2021) [7]. Version 4.2a of the database was chosen as the last release before the CAGI6 challenge deadline. For each predictor, we extracted the rank score of all dataset variants using SnpSift [45]. The rank score of a variant is its percentile, scaled between 0 and 1, among all variants in dbNSFP, with higher rank scores corresponding to more deleterious predictions. If a variant was assigned multiple rank scores (e.g. if the method makes separate predictions for each affected transcript), we took the highest rank score.

#### VARITY

Predictions from the VARITY class of models [38] (VARITY_R, VARITY_R_LOO, VARITY_ER, VARITY_ER_LOO), were added to dbNSFP v4.4a (released on May 6, 2023 after the challenge deadline). However, the VARITY models were trained on ClinVar and HGMD data released prior to the challenge deadline (F. Roth and J. Wu, personal communication, July 12, 2023). The same procedure described above was used to extract VARITY scores from dbNSFP v4.4a.

#### MutPred2

Predictions from MutPred2 [27] on the evaluation dataset were provided by the original authors (V. Pejaver, personal communication, August 25, 2023). Scores were provided per affected isoform, and the most pathogenic score was chosen.

#### PrimateAI-3D

Predictions from PrimateAI-3D [46] for most human missense variants were provided by Illumina. If a variant mapped to multiple genes, the most pathogenic score was chosen.

#### ESM-1b

ESM-1b [35] variant effect scores were computed for a ll p ossible amino acid changes in most proteins in the human proteome by Brandes *et al.* [47]. SnpSift was used to annotate each dataset variant with affected Ensembl transcripts and corresponding amino acid changes. Because ESM-1b scores are indexed by UniProt identifiers, we used the UniProt ID mapping service (https://www.uniprot.org/id-mapping) to convert Ensembl transcripts to UniProt IDs. If a variant mapped to multiple proteins, the most pathogenic score was chosen.

#### EVE

EVE [36] provides variant effect predictions in the form of VCF files for 2951 human proteins. The VCF files were downloaded on May 23, 2023 from https://evemodel.org/download/bulk. If a variant mapped to multiple proteins, the most pathogenic score was chosen.

#### AlphaMissense

We downloaded AlphaMissense [34] scores for variants in canonical isoforms and non-canonical isoforms on September 19, 2023 from https://console.cloud.google.com/storage/browser/dm_alphamissense. If a variant mapped to multiple isoforms, the most pathogenic score was chosen.

## Acknowledgements

This work was supported in part by the U.S. National Institutes of Health (NIH) awards U24HG007346 (S.E.B.), U41HG007346 (S.E.B.), R13HG006650 (S.E.B.), and U01HG012022 (P.R.). N.M.I. is a Chan Zuckerberg Biohub Investigator. We also thank the Belgian Fund for Scientific Research (F.R.S.–FNRS) and the Research Foundation Flanders (FWO) for financial support. The CAGI Annotate-All-Missense challenge was originally proposed by Sean Mooney.

## Supplementary Figures

**Fig. S1:**
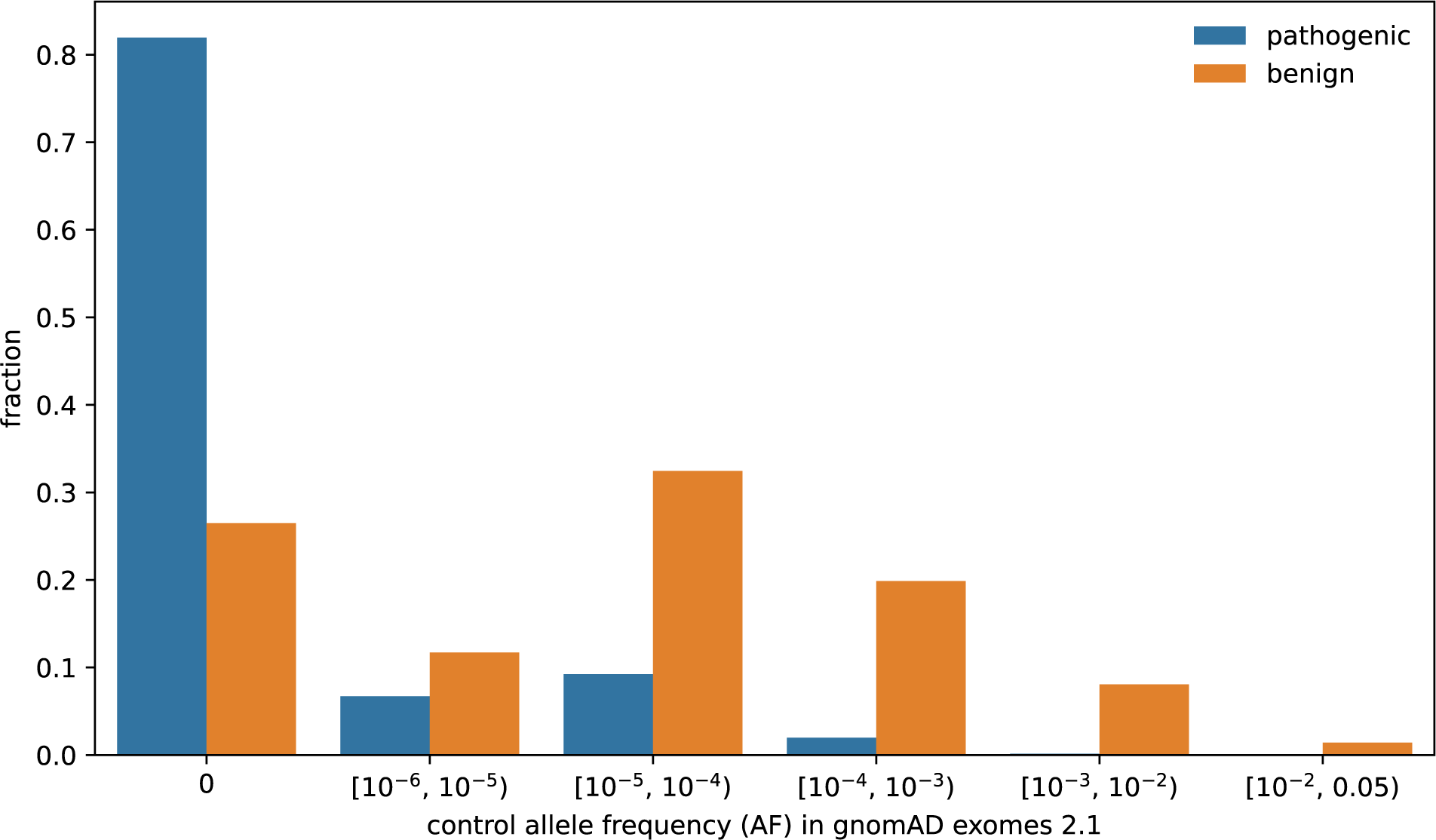
Allele frequency distributions of pathogenic and benign variants in the full evaluation dataset. Allele frequencies were obtained for all variants from the control cohort exomes in gnomAD v2.1.1 [13].

**Fig. S2:**
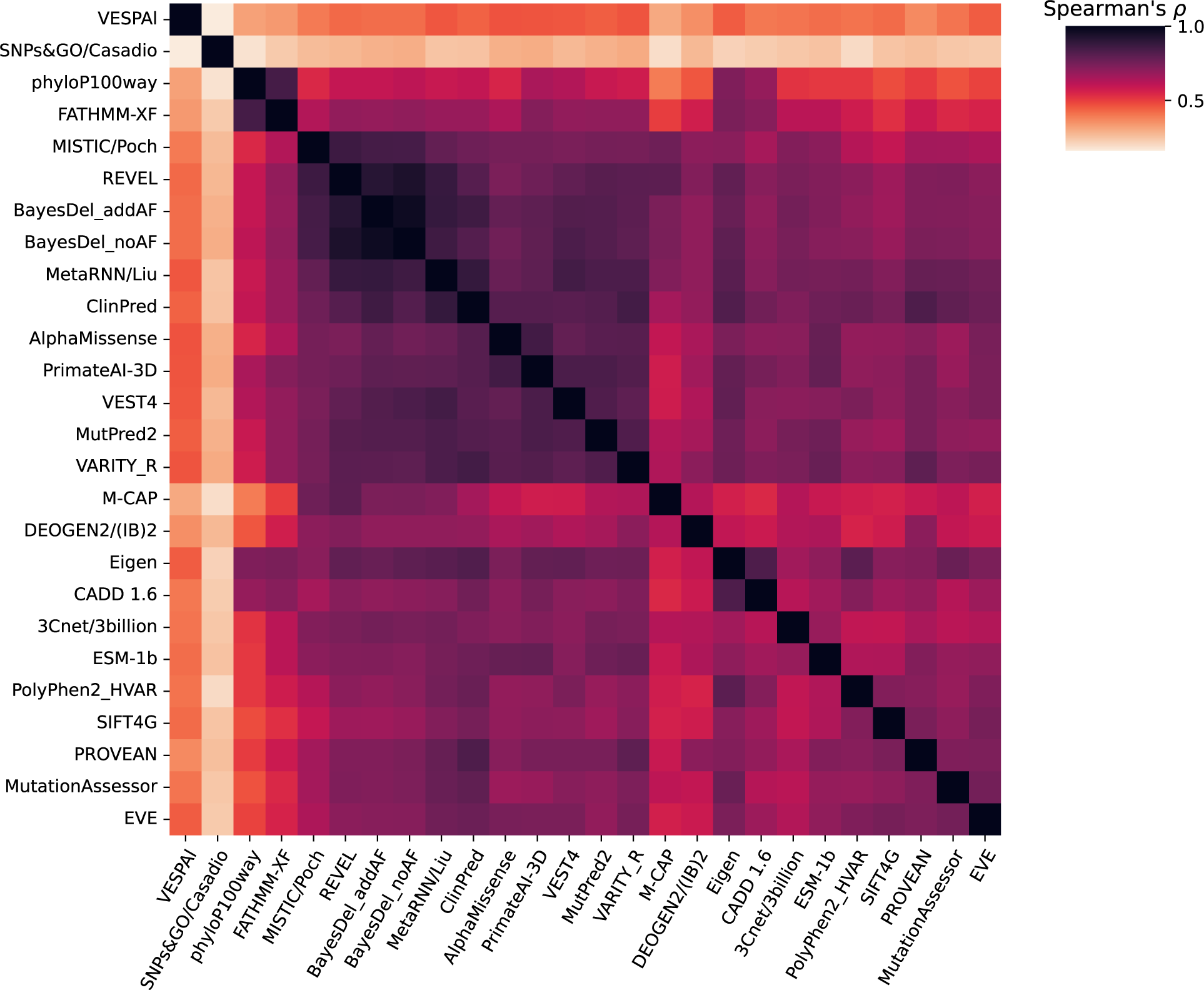
Correlations between the predictions of different tools. The heatmap shows the Spearman rank correlation coefficients between predictions computed on variants in the full evaluation dataset.

**Fig. S3:**
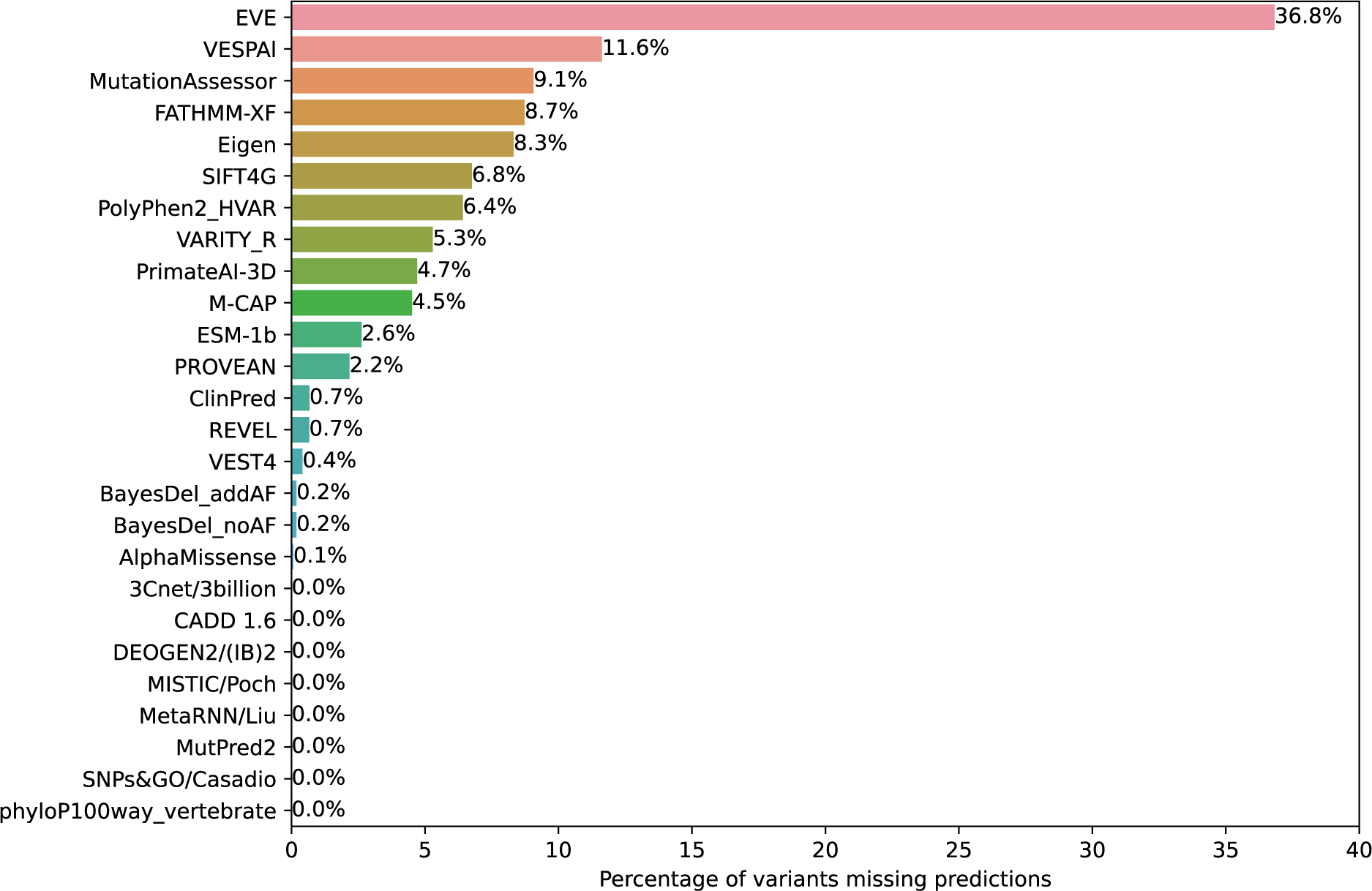
Missing predictions on evaluated variants. The percentage of variants within the full evaluation dataset for which predictions are not provided by each tool.

**Fig. S4:**
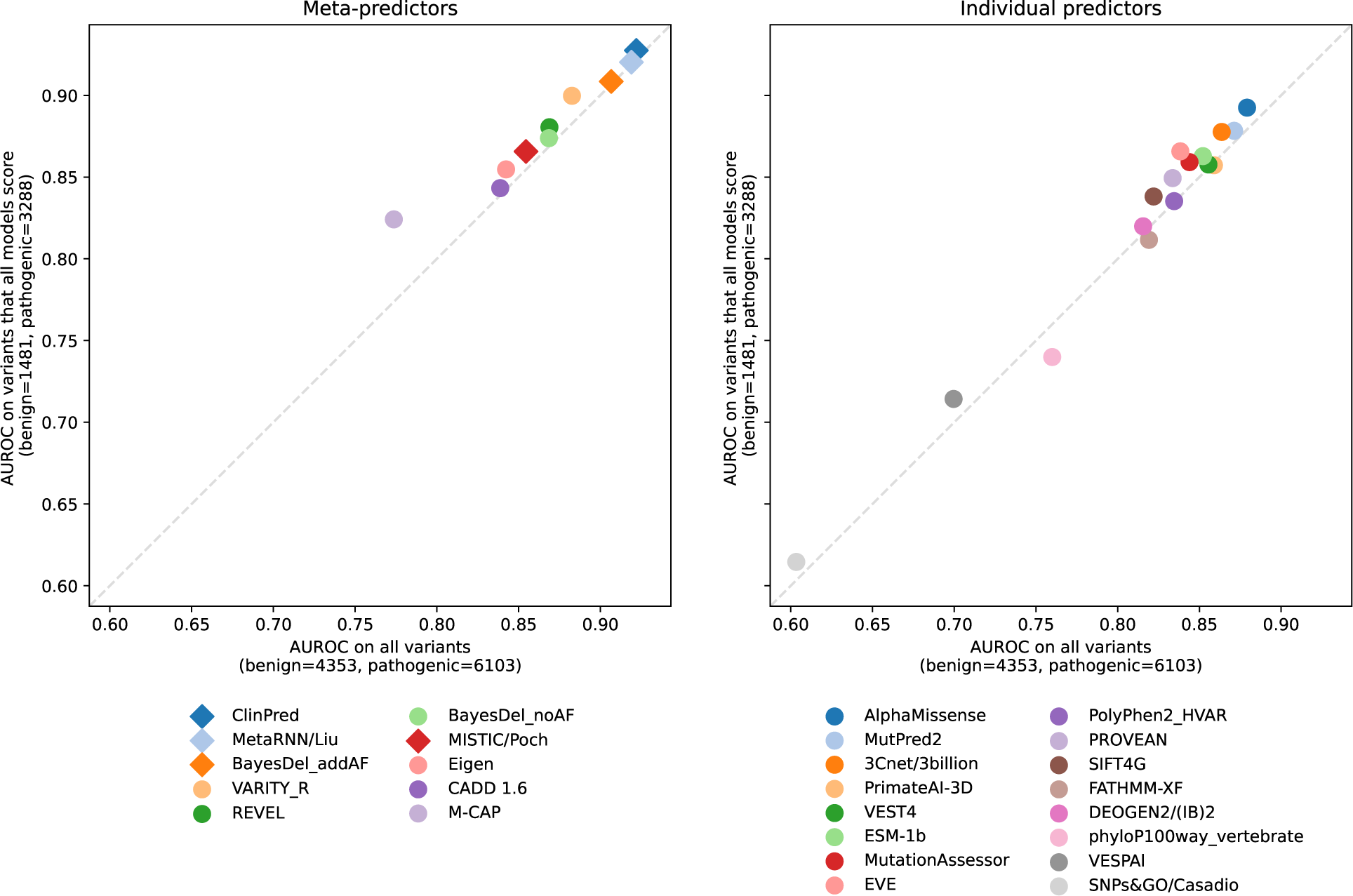
Performance on the subset of variants scored by all predictors. We compare AUROCs on the full set of variants (Fig. 1; x-axis) to AUROCs on the subset of variants scored by all predictors (y-axis) for meta-predictors (left) and individual predictors (right). Predictors marked by diamonds use allele frequency as a feature.

**Fig. S5:**
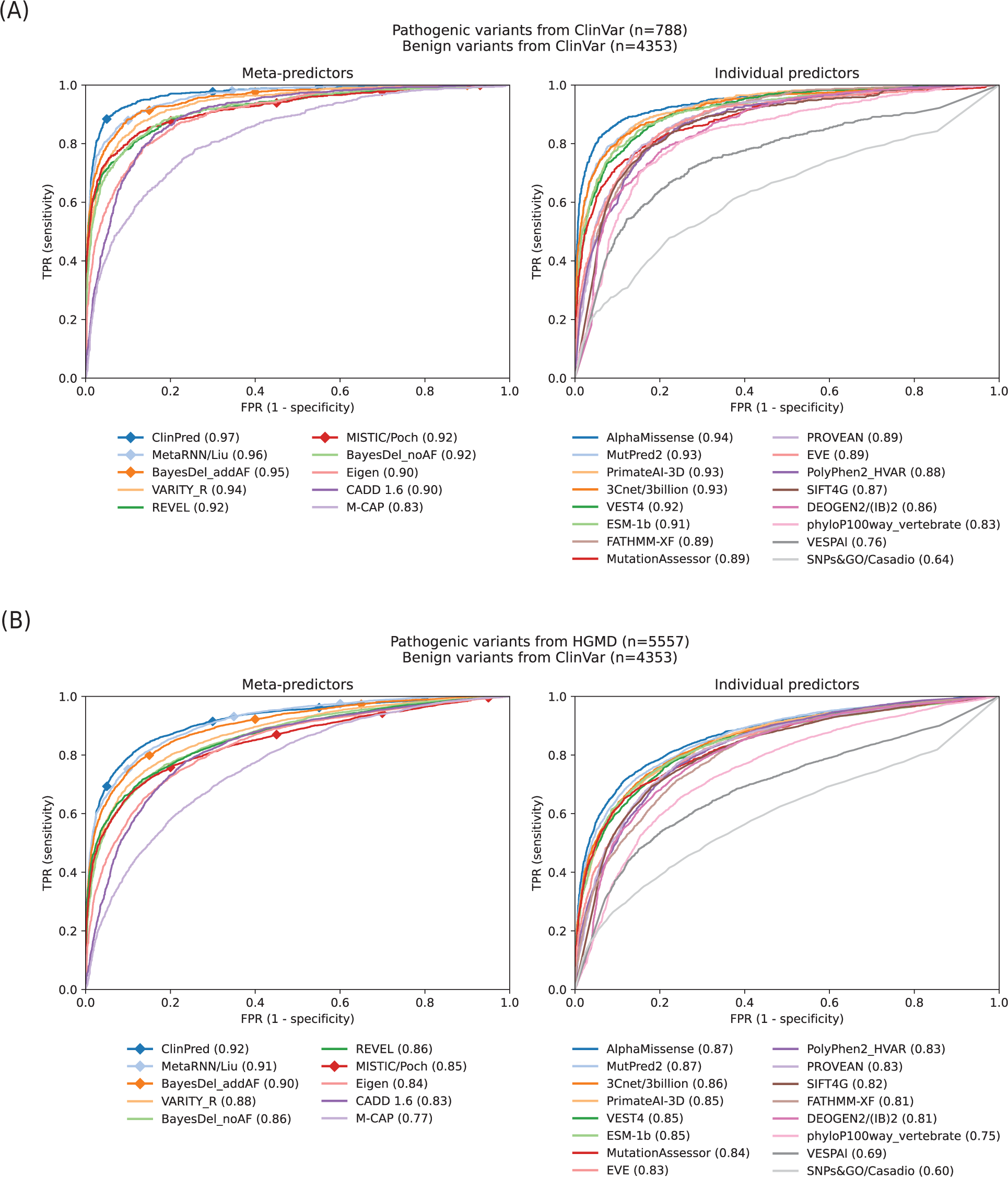
Performance when evaluating on only ClinVar or only HGMD pathogenic variants. We show the ROC curves and AUROCs for meta-predictors (left) and individual predictors (right) using pathogenic variants from either (A) ClinVar or (B) HGMD. Benign variants remain the same (from ClinVar) in both cases. Predictors marked by diamonds use allele frequency as a feature.

**Fig. S6:**
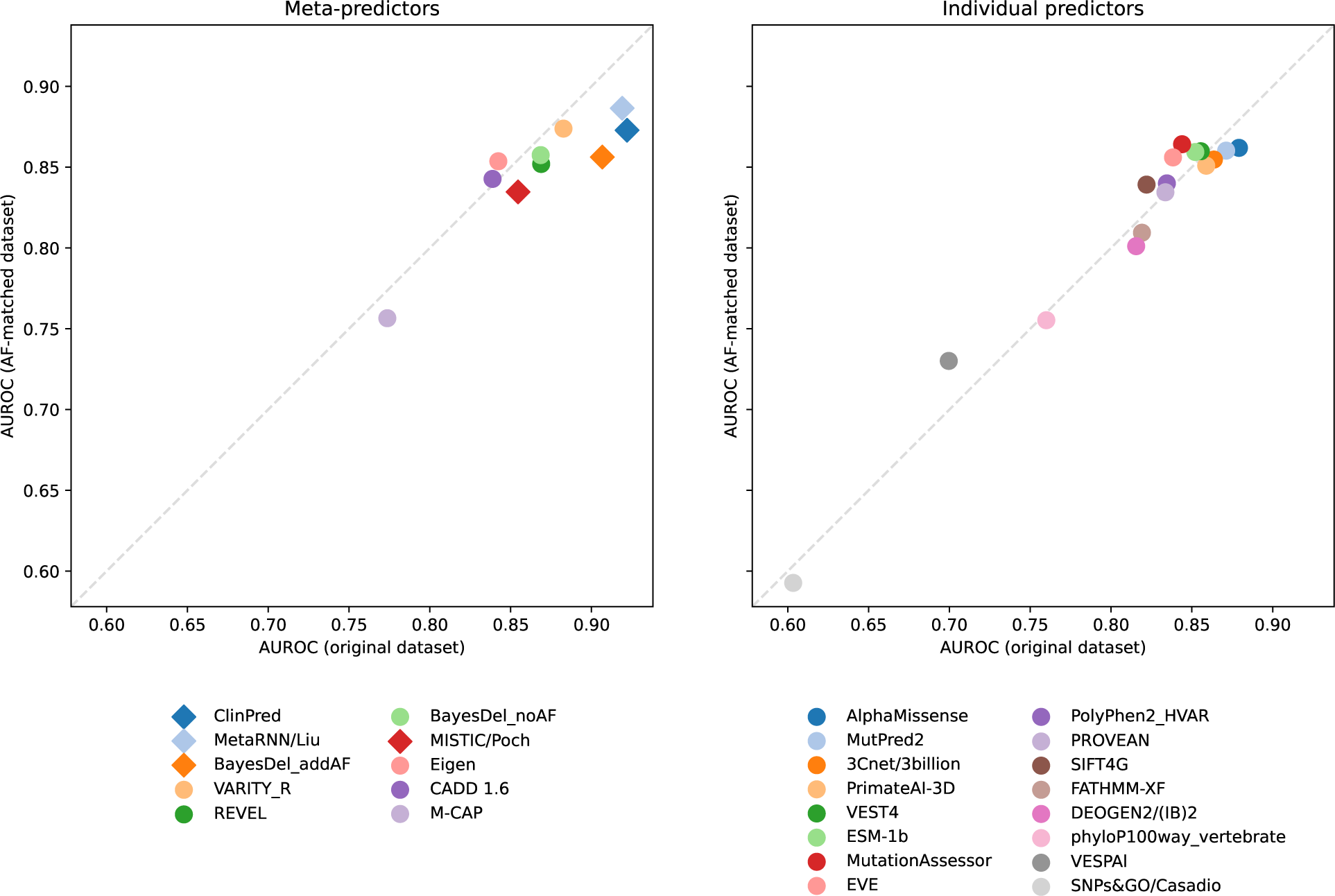
Performance on an allele frequency matched dataset. We created a subset of our evaluation dataset in which the allele frequency (AF) distributions of benign and pathogenic variants are matched. This matching was achieved by equalizing histogram bin counts in the log-transformed allele frequency space, using the Freedman-Diaconis rule [48] to determine bin widths. (See Fig. S1 to visualize the skew in the original dataset). We compare AUROCs on the full dataset (x-axis) to AUROCs on the allele frequency matched dataset (y-axis). Predictors marked by a diamond use allele frequency as a feature.

**Fig. S7:**
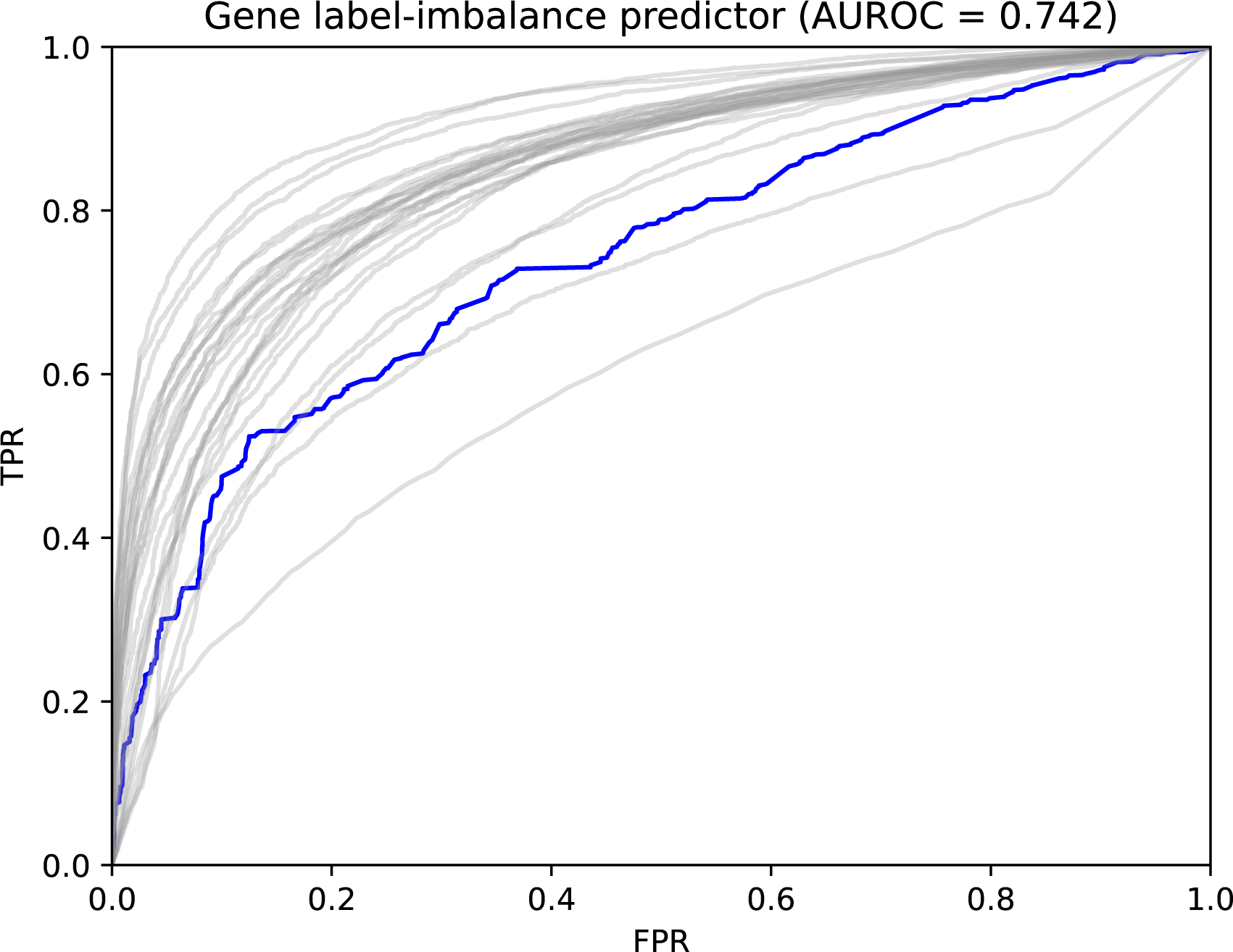
Performance of the gene label-imbalance baseline model. The blue ROC curve for the gene label-imbalance baseline model, which assigns the same score to all variants in a gene (equal to the fraction of ClinVar and HGMD missense variants that were labeled as pathogenic or DM before the cutoff date for our evaluation dataset), is overlayed on top of the light gray ROC curves for all assessed predictors on the full evaluation dataset.

**Fig. S8:**
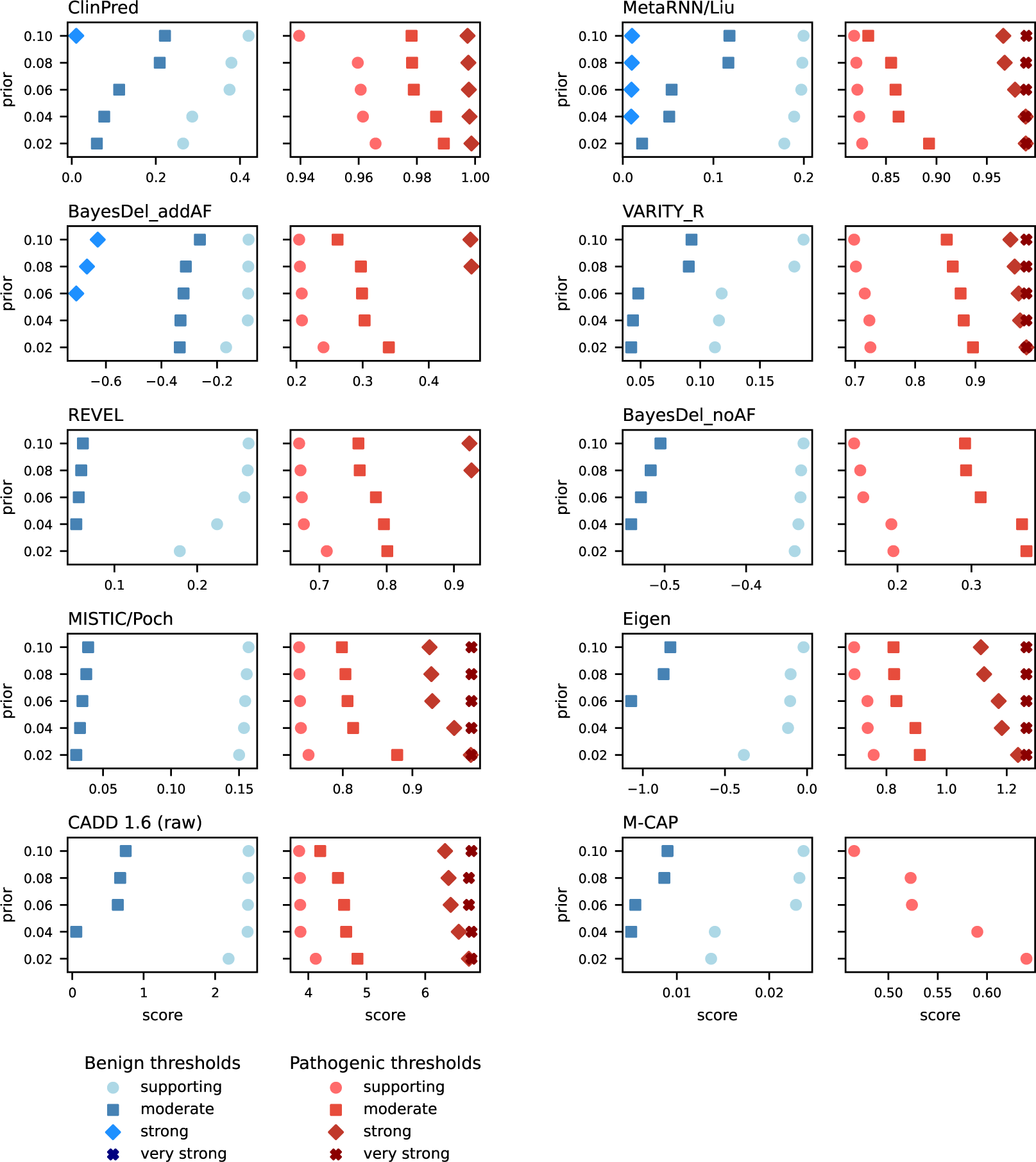
Meta-predictor score thresholds for benign and pathogenic evidence strengths at varying prior probabilities of pathogenicity. We determined the score thresholds that map to PP3 and BP4 evidence strengths by reimplementing the method proposed in [4] in Python. When computing the local posterior probability of pathogenicity (LP) for a given score *s*, our choice of interval around *s* differs slightly from that described in [4]. There, the authors recommend choosing an interval that contains the scores of at least 100 dataset variants and at least 3% of rare gnomAD variants. However, since the exact training sets of the predictors assessed here may overlap with gnomAD variants, we remove the gnomAD requirement and increase the number of dataset variants from 100 to 200 for smoothness of LP values.

**Fig. S9:**
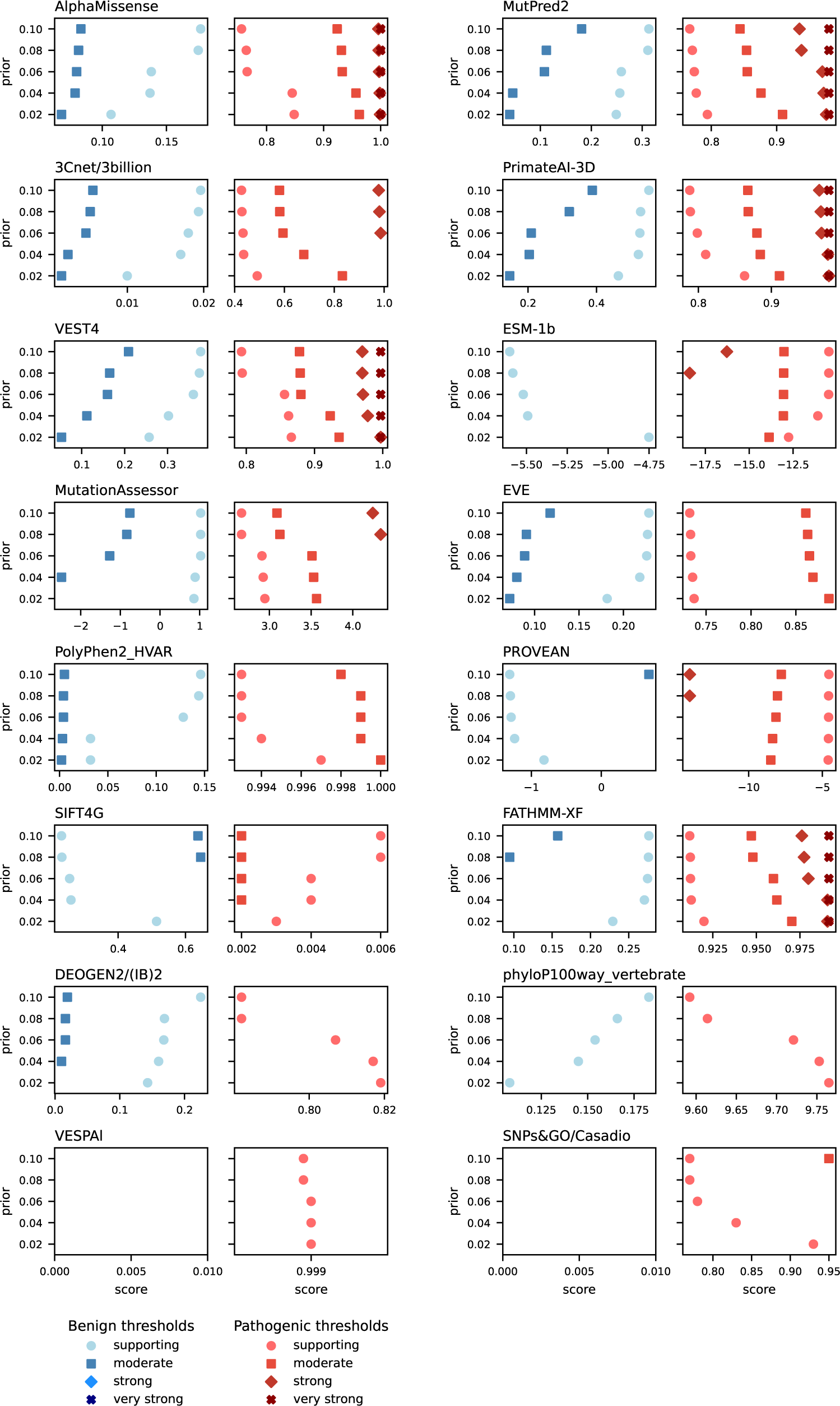
Individual predictor score thresholds for benign and pathogenic evidence strengths at varying prior probabilities of pathogenicity. We determined score thresholds as described in Fig. S8 above.

**Fig. S10:**
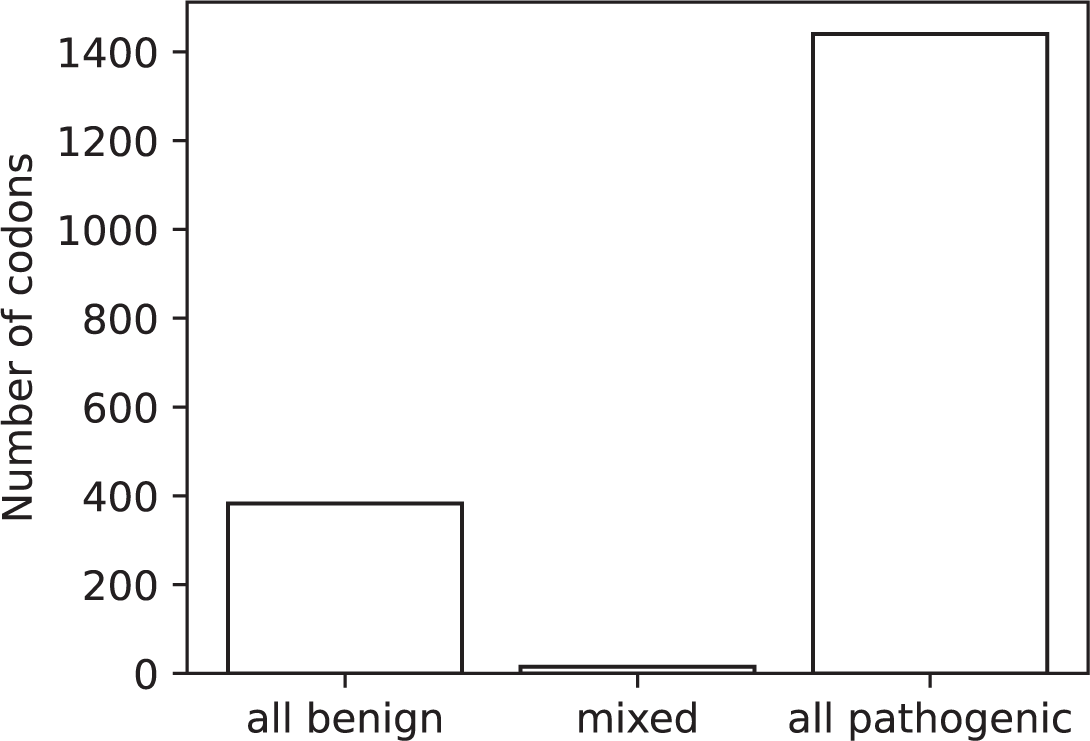
Codons with multiple missense variants. We used SnpEff [44] to determine the affected codon for all high-confidence missense variants in the April 4, 2023 version of ClinVar. We find that variants affecting the same codon are almost always classified in ClinVar as all benign or all pathogenic.

**Table S1:** Metrics for all predictors. For each of the 60 predictors evaluated, we show the AUROC from the full-dataset ROC curve (Fig. 1), as well as the high-specificity and high-sensitivity AUROCs (Fig. 2). We also include the standard deviation of each AUROC (calculated using 1000 bootstrap samples of the evaluation dataset) and the corresponding rank in each category. Predictors are ordered by their average rank across the three metrics. Note that this table combines all evaluated predictors, including meta-predictors and individual predictors, and predictors with and without allele frequency as a feature.

| predictor | full-dataset<br>AUROC±SD (rank) | high-specificity<br>AUROC±SD (rank) | high-sensitivity<br>AUROC±SD (rank) | average rank |
| --- | --- | --- | --- | --- |
| ClinPred [23] | 0.922 ± 0.003 (1) | 0.775 ± 0.008 (1) | 0.663 ± 0.009 (2) | 1.3 |
| MetaRNN/Liu [15] | 0.919 ± 0.002 (2) | 0.771 ± 0.007 (2) | 0.697 ± 0.007 (1) | 1.7 |
| BayesDel_addAF [21] | 0.907 ± 0.003 (3) | 0.757 ± 0.007 (3) | 0.653 ± 0.007 (3) | 3.0 |
| VARITY_R [38] | 0.883 ± 0.003 (4) | 0.716 ± 0.007 (7) | 0.621 ± 0.007 (5) | 5.3 |
| VARITY_R_LOO [38] | 0.883 ± 0.003 (5) | 0.716 ± 0.008 (6) | 0.621 ± 0.007 (6) | 5.7 |
| AlphaMissense [34] | 0.879 ± 0.003 (6) | 0.716 ± 0.007 (8) | 0.611 ± 0.006 (7) | 7.0 |
| VARITY_ER [38] | 0.875 ± 0.003 (7) | 0.713 ± 0.006 (10) | 0.610 ± 0.007 (8) | 8.3 |
| VARITY_ER_LOO [38] | 0.875 ± 0.003 (8) | 0.713 ± 0.007 (9) | 0.609 ± 0.007 (9) | 8.7 |
| MutPred2 [27] | 0.871 ± 0.003 (9) | 0.697 ± 0.007 (12) | 0.608 ± 0.007 (11) | 10.7 |
| REVEL [31] | 0.869 ± 0.003 (10) | 0.732 ± 0.006 (4) | 0.593 ± 0.006 (21) | 11.7 |
| BayesDel_noAF [21] | 0.869 ± 0.003 (11) | 0.702 ± 0.007 (11) | 0.602 ± 0.006 (13) | 11.7 |
| MutPred [49] | 0.860 ± 0.004 (14) | 0.667 ± 0.009 (19) | 0.622 ± 0.007 (4) | 12.3 |
| 3Cnet/3billion_model1 [14] | 0.864 ± 0.003 (12) | 0.689 ± 0.007 (15) | 0.600 ± 0.006 (14) | 13.7 |
| VEST4 [33] | 0.856 ± 0.004 (17) | 0.673 ± 0.007 (17) | 0.602 ± 0.006 (12) | 15.3 |
| 3Cnet/3billion_model2 [14] | 0.855 ± 0.004 (18) | 0.691 ± 0.007 (13) | 0.596 ± 0.006 (18) | 16.3 |
| 3Cnet/3billion_model4 [14] | 0.863 ± 0.004 (13) | 0.690 ± 0.008 (14) | 0.588 ± 0.006 (25) | 17.3 |
| PrimateAI-3D [37] | 0.859 ± 0.004 (16) | 0.667 ± 0.007 (20) | 0.596 ± 0.006 (17) | 17.7 |
| 3Cnet/3billion_model5 [14] | 0.859 ± 0.004 (15) | 0.658 ± 0.007 (24) | 0.600 ± 0.007 (15) | 18.0 |
| MISTIC/Poch [16] | 0.855 ± 0.004 (19) | 0.720 ± 0.006 (5) | 0.566 ± 0.004 (37) | 20.3 |
| 3Cnet/3billion_model6 [14] | 0.842 ± 0.004 (23) | 0.655 ± 0.007 (25) | 0.596 ± 0.006 (19) | 22.3 |
| 3Cnet/3billion_model3 [14] | 0.840 ± 0.004 (25) | 0.661 ± 0.007 (21) | 0.591 ± 0.006 (22) | 22.7 |
| PolyPhen2_HVAR [29] | 0.835 ± 0.004 (28) | 0.612 ± 0.006 (31) | 0.609 ± 0.006 (10) | 23.0 |
| MVP [50] | 0.842 ± 0.004 (24) | 0.659 ± 0.007 (22) | 0.589 ± 0.006 (24) | 23.3 |
| MutationAssessor [26] | 0.844 ± 0.004 (21) | 0.674 ± 0.007 (16) | 0.568 ± 0.006 (35) | 24.0 |
| ESM-1b [35] | 0.852 ± 0.004 (20) | 0.668 ± 0.007 (18) | 0.565 ± 0.005 (38) | 25.3 |
| Eigen [24] | 0.842 ± 0.004 (22) | 0.652 ± 0.007 (26) | 0.579 ± 0.006 (28) | 25.3 |
| EVE [36] | 0.838 ± 0.005 (27) | 0.633 ± 0.009 (28) | 0.589 ± 0.007 (23) | 26.0 |
| DEOGEN2 [19] | 0.832 ± 0.004 (30) | 0.658 ± 0.006 (23) | 0.577 ± 0.005 (32) | 28.3 |
| FATHMM-XF [25] | 0.819 ± 0.004 (33) | 0.607 ± 0.006 (32) | 0.595 ± 0.006 (20) | 28.3 |
| CADD 1.6 [22] | 0.839 ± 0.004 (26) | 0.597 ± 0.006 (34) | 0.587 ± 0.005 (26) | 28.7 |
| PROVEAN [20] | 0.834 ± 0.004 (29) | 0.603 ± 0.007 (33) | 0.578 ± 0.006 (29) | 30.3 |
| PolyPhen2_HDIV [29] | 0.812 ± 0.004 (36) | 0.558 ± 0.004 (41) | 0.598 ± 0.006 (16) | 31.0 |
| Eigen-PC [24] | 0.821 ± 0.004 (32) | 0.622 ± 0.007 (30) | 0.568 ± 0.005 (36) | 32.7 |
| MetaLR [51] | 0.798 ± 0.004 (39) | 0.629 ± 0.005 (29) | 0.578 ± 0.005 (31) | 33.0 |
| SIFT4G [32] | 0.822 ± 0.004 (31) | 0.572 ± 0.007 (38) | 0.570 ± 0.005 (34) | 34.3 |
| DEOGEN2/(IB)2 [19] | 0.816 ± 0.004 (34) | 0.544 ± 0.007 (46) | 0.584 ± 0.006 (27) | 35.7 |
| M-CAP [28] | 0.774 ± 0.005 (40) | 0.573 ± 0.005 (37) | 0.575 ± 0.004 (33) | 36.7 |
| MetaSVM [51] | 0.813 ± 0.004 (35) | 0.637 ± 0.006 (27) | 0.531 ± 0.004 (53) | 38.3 |
| SIFT [52] | 0.802 ± 0.004 (38) | 0.565 ± 0.005 (40) | 0.561 ± 0.005 (40) | 39.3 |
| SNPMuSiC/(IB)2 [53] | 0.804 ± 0.004 (37) | 0.567 ± 0.004 (39) | 0.551 ± 0.003 (44) | 40.0 |
| LIST-S2 [54] | 0.756 ± 0.005 (43) | 0.580 ± 0.005 (35) | 0.548 ± 0.004 (47) | 41.7 |
| DANN [55] | 0.743 ± 0.005 (44) | 0.517 ± 0.003 (52) | 0.578 ± 0.005 (30) | 42.0 |
| MutationTaster [56] | 0.757 ± 0.005 (42) | 0.522 ± 0.001 (48) | 0.562 ± 0.005 (39) | 43.0 |
| PrimateAI [46] | 0.734 ± 0.005 (46) | 0.549 ± 0.005 (43) | 0.550 ± 0.004 (45) | 44.7 |
| phyloP100way_vertebrate [30] | 0.760 ± 0.005 (41) | 0.539 ± 0.004 (47) | 0.543 ± 0.003 (48) | 45.3 |
| LRT [57] | 0.737 ± 0.005 (45) | 0.521 ± 0.001 (49) | 0.553 ± 0.005 (42) | 45.3 |
| FATHMM-MKL [58] | 0.733 ± 0.005 (47) | 0.521 ± 0.003 (50) | 0.559 ± 0.004 (41) | 46.0 |
| MPC [59] | 0.712 ± 0.005 (48) | 0.577 ± 0.005 (36) | 0.525 ± 0.004 (55) | 46.3 |
| FATHMM [60] | 0.677 ± 0.005 (51) | 0.551 ± 0.004 (42) | 0.532 ± 0.003 (51) | 48.0 |
| phastCons100way_vertebrate [61] | 0.674 ± 0.005 (52) | 0.508 ± 0.000 (55) | 0.552 ± 0.004 (43) | 50.0 |
| VESPAI [18] | 0.700 ± 0.005 (49) | 0.545 ± 0.004 (45) | 0.506 ± 0.001 (58) | 50.7 |
| SiPhy_29way_logOdds [62] | 0.683 ± 0.005 (50) | 0.507 ± 0.002 (56) | 0.549 ± 0.004 (46) | 50.7 |
| GERP++_RS [63] | 0.668 ± 0.006 (53) | 0.509 ± 0.002 (54) | 0.538 ± 0.004 (49) | 52.0 |
| phyloP30way_mammalian [30] | 0.627 ± 0.005 (55) | 0.509 ± 0.001 (53) | 0.536 ± 0.003 (50) | 52.7 |
| SNPs&GO/Casadio [17] | 0.603 ± 0.005 (56) | 0.548 ± 0.004 (44) | 0.498 ± 0.000 (60) | 53.3 |
| GenoCanyon [64] | 0.645 ± 0.005 (54) | 0.520 ± 0.002 (51) | 0.515 ± 0.002 (57) | 54.0 |
| phyloP17way_primate [30] | 0.589 ± 0.006 (58) | 0.506 ± 0.001 (57) | 0.531 ± 0.003 (52) | 55.7 |
| phastCons30way_mammalian [61] | 0.599 ± 0.006 (57) | 0.506 ± 0.001 (58) | 0.531 ± 0.003 (54) | 56.3 |
| phastCons17way_primate [61] | 0.574 ± 0.006 (59) | 0.503 ± 0.001 (59) | 0.521 ± 0.002 (56) | 58.0 |
| bStatistic [65] | 0.497 ± 0.006 (60) | 0.499 ± 0.001 (60) | 0.499 ± 0.001 (59) | 59.7 |

